# Inter-organ regulation of *Drosophila* intestinal stem cell proliferation by a hybrid organ boundary zone

**DOI:** 10.1101/152074

**Authors:** Jessica K. Sawyer, Erez Cohen, Donald T. Fox

**Affiliations:** Department of Pharmacology & Cancer Biology, Duke University Medical Center, DUMC Box 3813, Durham, NC 27710; Department of Cell Biology Duke University Medical Center, DUMC Box 3813, Durham, NC 27710; Regeneration Next, Duke University Medical Center, DUMC Box 3813, Durham, NC 27710

**Keywords:** Drosophila, hindgut, midgut, organ boundary, intestinal stem cell

## Abstract

Gene expression at the *Drosophila* midgut-hindgut boundary is a hybrid of both organs. Hybrid cells repress stem cell division, but boundary injury activates stem cell division through inter-organ JAK-STAT signaling.

**ABSTRACT:** The molecular identities and regulation of cells at inter-organ boundaries are often unclear, despite the increasingly appreciated role of organ boundaries in disease. Using *Drosophila* as a model, here we show that a specific population of adult midgut organ boundary intestinal stem cells (OB-ISCs) is regulated by the neighboring hindgut, a developmentally distinct organ. This distinct OB-ISCs control is due to proximity to a specialized transition zone between the endodermal midgut and ectodermal hindgut that shares molecular signatures of both organs, which we term the hybrid zone (HZ). During homeostasis, proximity to the HZ restrains OB-ISC proliferation. However, injury to the adult HZ/hindgut drives up-regulation of *upaired-3* cytokine and OB-ISC hyperplasia. If HZ disruption is severe, hyperplastic OB-ISCs expand across the inter-organ boundary. Our data suggest that inter-organ signaling plays an important role in controlling OB-ISCs in homeostasis and injury repair, which is likely critical in prevention of disease.

## INTRODUCTION

Inter-organ boundaries, such as those in the intestine, provide an opportunity to understand an emerging area of stem cell research: signaling to stem cells from a neighboring tissue. Stem cells at inter-organ boundaries are near cells from a functionally distinct organ. Therefore, such stem cells may reside in a very different tissue environment than other seemingly identical stem cells elsewhere in the same organ. Very little is known about stem cell control at intestinal organ boundaries, yet hyper-proliferation of putative injury activated stem cells and altered cell fate at the human esophagus/stomach boundary is linked to Barrett’s esophagus, a condition that increases cancer risk (Badreddine and Wang, 2010; Hvid-Jensen et al., 2011; San Roman and Shivdasani, 2011). New understanding of the complex inter-organ stem cell control at intestinal organ boundaries could be aided by development of a simple, genetically tractable model tissue.

Insect intestinal boundaries are an excellent candidate to model inter-organ stem cell control. At both the foregut/midgut and midgut/hindgut organ boundaries are cell populations classically defined as imaginal rings, which are enriched for Wnt ligands (Bodenstein, 1950; Fox and Spradling, 2009; Robertson, 1936; Takashima et al., 2008; Tian et al., 2016). The boundary between the endodermal midgut (small intestine) and ectodermal hindgut (large intestine) demarcates a sharp contrast in intestinal cell architecture and cell turnover rates. The adult *Drosophila* midgut, which is rich in microvilli (Fig1A’), turns over weekly under rich diet conditions. The adult midgut is comprised of multipotent intestinal stem cell (ISCs) and their progeny: the secretory enteroendocrine cells (ees) and enteroblasts (EBs), which differentiate into polyploid absorptive enterocytes (ECs) (Guo and Ohlstein, 2015; Micchelli and Perrimon, 2006; Ohlstein and Spradling, 2006, 2007; Zeng and Hou, 2015). This proliferative organ responds to tissue loss via compensatory stem cell proliferation (Apidianakis et al., 2009; Biteau et al., 2008; Buchon et al., 2009a; Buchon et al., 2009b; Chatterjee and Ip, 2009; Jiang et al., 2009). Adjacent to the posterior midgut is the pyloric sphincter of the *Drosophila* hindgut (Fig1A, in red), the functional analog of the mammalian ileocecal sphincter, which lacks microvilli but is chitin-rich (Fig1A’). Under normal homeostatic conditions, the pylorus is a quiescent diploid tissue as it has negligible levels of cell cycle activity (Fox and Spradling, 2009). Following injury, the pylorus does re-enter the cell cycle, but instead of cell division it undergoes compensatory cellular hypertrophy and wound-induced endocycles, which replicate the DNA without cell division, generating polyploid pyloric cells (Losick et al., 2013). We previously proposed that injury-responsive stem cells were in the Wnt/Wingless (Wg) positive imaginal ring region (hereafter: Wg ring), but it remained possible that these cells were adjacent to the Wg ring (Fox and Spradling, 2009). The distinct cell types of the midgut/hindgut boundary and their potential role in the homeostasis/tissue repair program of each organ remained to be determined.

**Figure 1.**
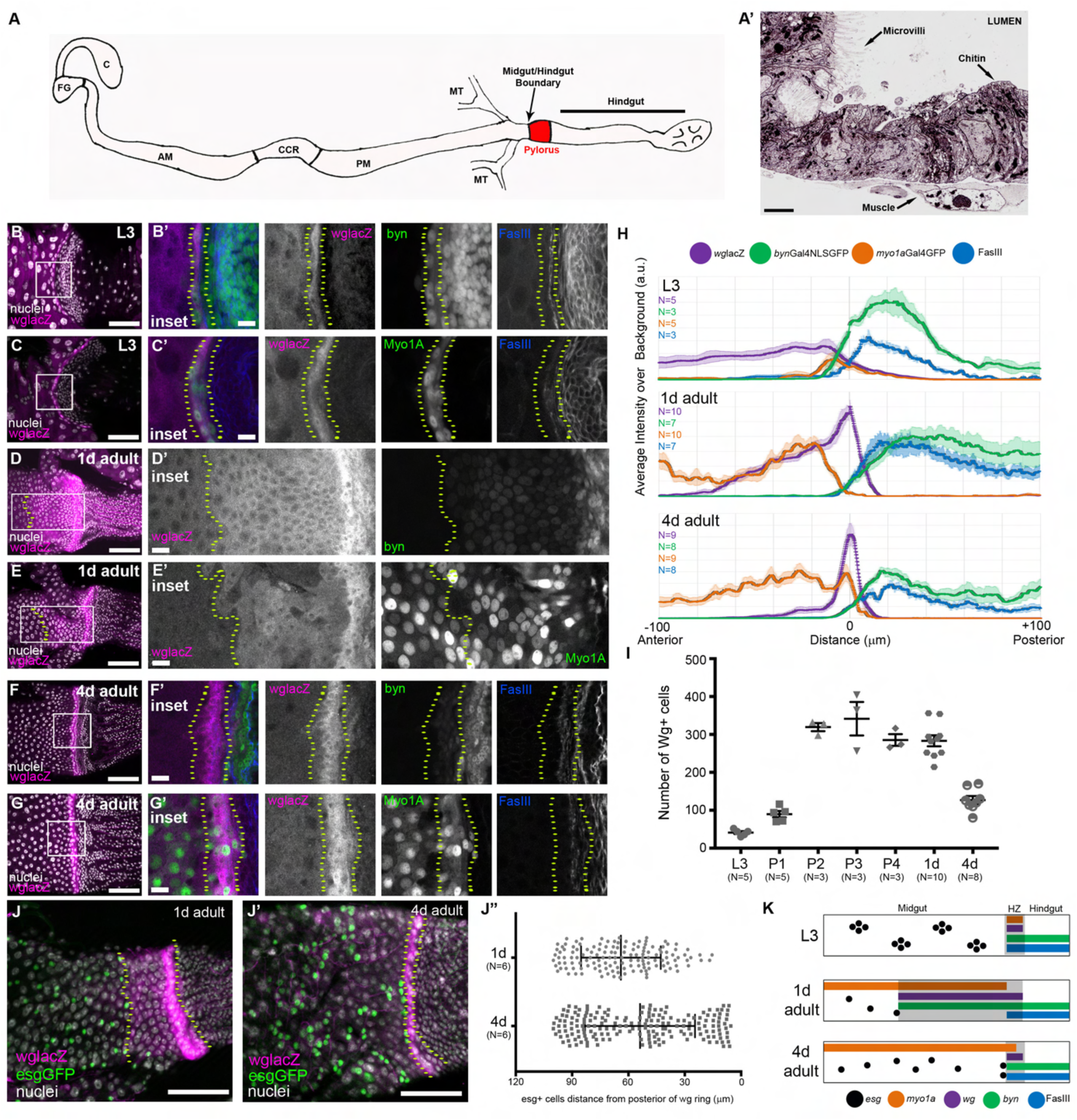
The Wg ring is a dynamic, hybrid zone adjacent to midgut ISCs/EBs. (A) Schematic of the *Drosophila* intestine; foregut (FG), crop (C), anterior midgut (AM), copper cell region (CCR), posterior midgut (PM), malpighian tubules (MT). (A’) Electron micrograph illustrating the phenotypic differences at the midgut/hindgut boundary. There is a clear transition from cells with microvilli to cells that lack microvilli, but are chitin-rich, scale=2μm. (B-I) The HZ is present throughout development: L3 (B-C), 1d adult (D-E), 4d adult (F-G), scale=50μm and 10μm. (H) Line profiles of midgut, HZ, and hindgut markers. Data represent mean ± SEM. (I) The number of *wg*+ cells over time, L3 to adulthood. Data represent mean ± SEM. (J) *esg*+ cells stay at the anterior edge of *wg* expression domain, scale=50μm. Data represent mean ± SEM of all *esg*+ from 6 animals. (K) Model of HZ during development. Genotypes and markers indicated in panels, yellow dotted lines indicate the HZ.

Here, we characterize the cellular architecture and function of the *Drosophila* midgut/hindgut boundary. In doing so, we show that the adult Wg imaginal ring is a transition zone that shows hybrid (both midgut and hindgut) gene expression. The function of this hybrid zone (HZ) is to repress proliferation of organ-boundary intestinal stem cells (OB-ISCs) located in the midgut, which reside immediately adjacent to the anterior boundary of the HZ. Relative to most other posterior midgut ISCs, OB-ISCs exhibit low cell cycle rates and are resistant to *Notch*-mediated stem cell tumor formation. When injury locally disrupts the hybrid zone, OB-ISC hyperplasia is induced. Under conditions of severe injury, hyperplastic OB-ISCs often cross the midgut/hindgut boundary through disrupted segments of the HZ. This hyperplasia coincides with injury-induced release of the JAK-SAT ligand Unpaired-3 (Upd3) from the hindgut and HZ, which is sufficient to induce OB-ISC proliferation. Our results identify a HZ as a strategy for locally regulating signaling between adjacent organs. We also find that interactions between organs at a boundary can profoundly influence stem cell activity. Thus, use of a HZ between different organs can preserve organ integrity and maintain cell fate between two distinct, yet adjacent, organs.

## RESULTS

### The Wg ring is a dynamic, hybrid zone adjacent to midgut ISCs/EBs

We previously identified putative stem cell activity at the injured adult *Drosophila* midgut/hindgut boundary (Fox and Spradling, 2009). Given the complexity of cell types in this region, we next characterized each epithelial population in the midgut/hindgut boundary at single-cell resolution. A ring of epithelial cells that strongly expresses the Wnt ligand *wg* (hereafter Wg ring) resides directly at the midgut/hindgut boundary (Fox and Spradling, 2009; Takashima et al., 2008; Takashima and Murakami, 2001; Tian et al., 2016). To determine whether the cells in the Wg ring possess molecular characteristics of hindgut or midgut, we used a *lacZ* trap of the *wg* gene (*wglacZ*) to mark this population and then examined two well-known markers each of both the midgut and hindgut. First, we examined animals during late 3^rd^ instar larval development (L3). The Wg ring during L3 is comprised of ~40 cells that also express two hindgut markers: the pan-hindgut transcription factor *brachyenteron* (*byn*) and the cell adhesion molecule Fasciclin III (FasIII) (Fig.1B-B’,H,I). Anterior to the Wg ring, we did not detect the expression of *byn* and FasIII is weakly accumulated. Additionally, the same *byn*+FasIII+ cells of the L3 Wg ring cells also express *Myosin 31DF* (*Myo1a*), a marker of midgut differentiated enterocytes, as previously reported (Gonzalez-Morales et al., 2015), Fig1C-C’,H,I). We further found that a second midgut enterocyte marker, nubbin/POU domain protein 1 (Pdm1), co-localizes with *Myo1a* in the L3 Wg ring (SupplFig1A-A”). Together, these results suggest that just prior to metamorphosis, cells of the Wg ring simultaneously express not only multiple hindgut markers but also well-established markers of midgut enterocytes.

During metamorphosis, much of the intestinal epithelium undergoes histolysis and is reformed, while the region around the Wg ring remains intact (Fox and Spradling, 2009; Robertson, 1936; Takashima et al., 2008). We next examined how cell fate and number in the Wg ring are altered during metamorphosis. At two distinct stages of pupal development (day 1 and 3 post-puparium formation), *wg* and *byn* expression in the Wg ring remain tightly correlated (SupplFig1C-C”’,E-E”’). We also find *Myo1a* expression in some *wg*+*byn*+ cells, but not all (SupplFig1D-D’”,F-F”’). The Wg ring remains present throughout intestinal remodeling and continues to express markers of both the midgut and hindgut. During pupation, the size of the Wg domain expands by ~7 fold in cell number (Fig1I). As adulthood begins (1d), the Wg domain can be sub-divided into two regions. In the anterior region, low level *wg*+ cells express *byn* and *Myo1a,* and to a lesser extent FasIII (Fig1D-E’,H). Closer to the hindgut, we detected strong *byn* expression but very low levels of *Myo1a* in high *wg*+ expressing cells (Fig1D-E’). In mature adults (4d, after gut remodeling is complete) the Wg domain shrinks to ~125 cells (Fig1F-G’,H,I). We propose that this is due to expression changes, as we did not see evidence of cell death in early adulthood (SupplFig1G-G”, see Fig4B-B’ for positive control). As the Wg domain recedes, we observe the emergence of adult midgut ISCs and their immediate EB daughters, marked by expression of the transcription factor *escargot* (*esg*), in the posterior midgut next to the midgut/hindgut boundary (Fig1J-J”). This finding is consistent with previous reports that ISCs may migrate to the posterior midgut (Takashima et al., 2016; Takashima et al., 2013) and suggests that the extent of the Wg domain determines the posterior position of midgut ISCs/EBs.

Our data from larval and pupal stages suggested that many *wg*+ cells may be uncommitted to a particular organ identity during development. We next investigated whether the mature adult Wg ring retains a mixed organ identity or acquires a specific organ fate. Our marker expression analysis indicated that the adult Wg ring continues to co-express *byn*, FasIII, *Myo1a*, and Pdm1 and retain the hybrid organ characteristics (Fig1F-G’,H,SupplFig1B-B”’). Thus, the Wg domain is dynamic in size from late larva (L3) to mature adult. Further, at all stages examined, the Wg ring domain contains cells that express markers of both the midgut and hindgut (Fig1K). Due to the persistent dual-organ gene expression in this midgut/hindgut transition zone, we term this region the hybrid zone (HZ).

### The adult HZ and adjacent posterior midgut arise from distinct progenitor pools

Previous studies suggested that during metamorphosis, the epithelial region containing the adult HZ and a portion of the posterior midgut arises from *byn+,FasIII+* cells in the larval hindgut pylorus that trans-differentiate into *byn-,FasIII-, Myo1A*+,Pdm1+ enterocytes (Fig2A; Takashima et al., 2008; Takashima et al., 2013). However, these studies used population-wide lineage tracing approaches, which have the caveat that they cannot distinguish subpopulations of cells expressing a given promoter (Fox et al., 2008). Additionally, our finding that a hybrid midgut/hindgut population persists throughout metamorphosis (SupplFig1C and E) suggests that the HZ and adjacent posterior midgut likely arise from distinct progenitor pools, rather than from trans-differentiation of pyloric cells (Takashima et al., 2013). To test whether the HZ and adjacent posterior midgut share a common progenitor with the pylorus at the onset of metamorphosis, we followed the lineage of individually labeled cells using *hsFLP;; actin-FLPout-lacZ*. In addition to providing unbiased, random labeling, we observed little to no background labeling with this system (SupplFig2A-B”). This low background labeling enabled us to avoid confusing our labeling with non-induced labeling at unspecified times in development/adulthood. To determine how the larval HZ develops into the adult HZ during metamorphosis, we induced a single low level heat shock in wandering L3 animals and examined the clones in resulting four day old adults (prior to significant turnover of adult midgut cells, see Methods).

**Figure 2.**
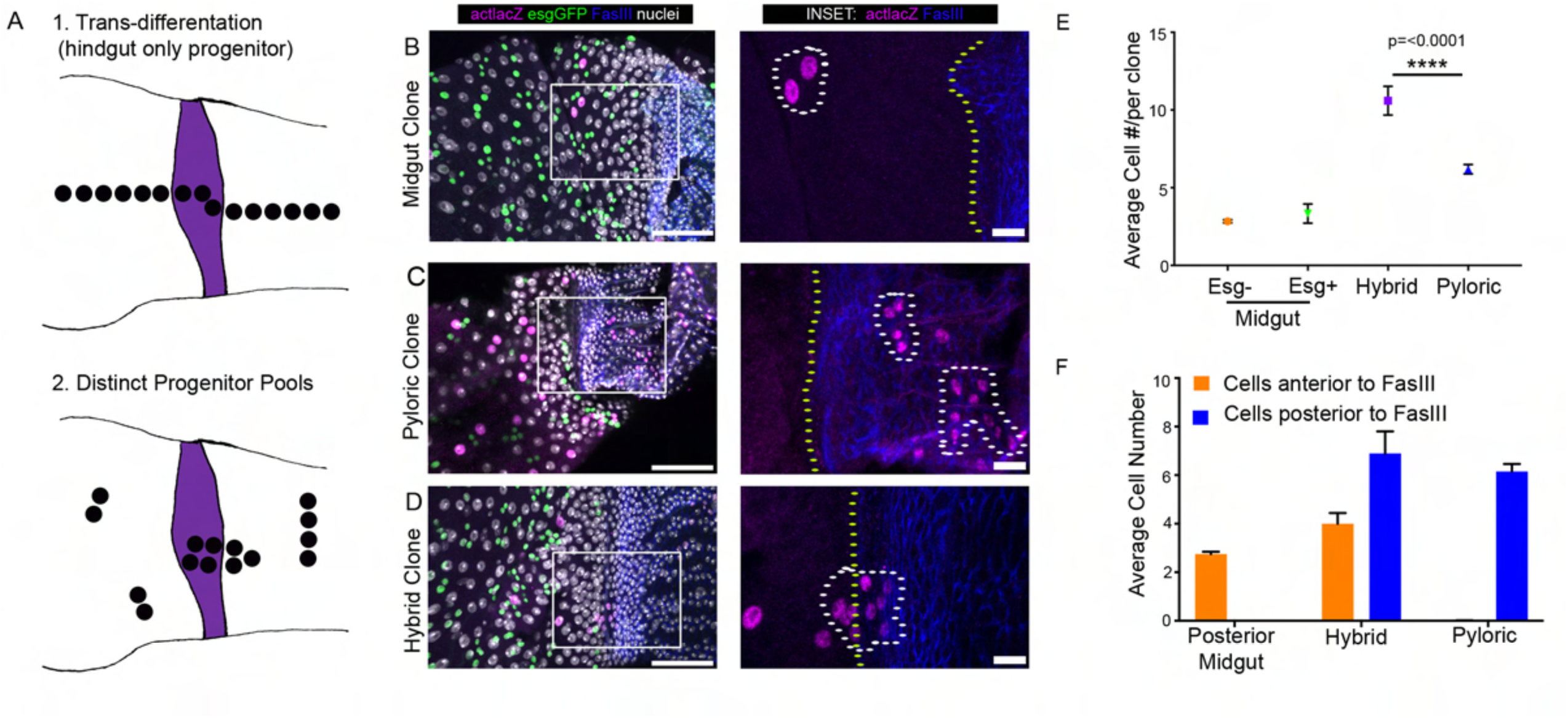
The adult HZ and adjacent posterior midgut arise from distinct progenitor pools. (A) Model comparing expected clonal results for trans-differentiation vs. distinct progenitor pools. (B-D) Examples of clone types observed: midgut (B), pyloric (C), hybrid (D), scale=50μm and 10μm. (E) Hybrid clones are significantly larger than midgut and pyloric clones. Data represent mean ± SEM. (F) Hybrid clones cross the FasIII border. Data represent mean ± SEM. Genotypes and markers indicated in panels, yellow dotted lines indicate the HZ, white dotted lines indicate clones.

Our lineage results from 316 independent clones did not reveal any evidence of continuous clones that encompass the pylorus, HZ, and adjacent posterior midgut. Rather, based on clone location and size, we observed three distinct epithelial clonal patterns in the region from the posterior midgut to the adult hindgut pylorus (Fig2A-D). The first pattern we observed consist of clones not in the HZ, but in the nearby posterior midgut (55% of all clones, avg size 2.9±0.1, Fig2B,E,SupplFig2C). The vast majority (98%) of these clones consisted only of polyploid midgut enterocytes. A small subset of these midgut clones (2%) contained ISCs/EBs (*esg*+). We never observed overlap between *esg*+ clones and the HZ, suggesting *esg+* ISCs/EBs do not generate the majority of the posterior-most midgut region or the adult HZ, in agreement with previous findings (Takashima et al., 2013). The second class of clones that we observed resided entirely within the pylorus (hindgut) and not in the HZ (38% of all clones, avg size 6.2±0.3, Fig2C,E,SupplFig2C). Finally, we observed a unique class of HZ-localized clones (7% of all clones, avg size 10.6±0.9, Fig2D,E,SupplFig2C).

HZ clones were distinct from midgut and pyloric clones in multiple ways. In addition to being significantly larger than midgut or pyloric clones (Fig2E), only this class of clones overlaps the midgut/hindgut boundary as defined by FasIII (Fig2D,F). Typically, HZ clones have two more cells posterior to the FasIII boundary than anterior (6 cells posterior vs 4 anterior, Fig2E). By co-imaging with a marker of the HZ, we found that HZ clones are either contained entirely within the HZ or extend up to 6-7 cell diameters into the anterior pylorus (SupplFig2D). Further, none of the HZ clones (0/20) contain *esg*+ ISCs/EBs. This finding reinforces the model that the HZ does not arise from these midgut cells and also shows that *esg*+ cells and HZ cells have distinct developmental origins. Taken together, these clonal data rule out the idea that a common hindgut progenitor generates the adult pylorus, HZ, and adjacent posterior midgut during metamorphosis. It remains possible that very early in development (prior to the 3^rd^ instar) these three distinct adult populations share a common progenitor. Further, given both our clonal data the expansive nature of the HZ during metamorphosis (FigSuppl1), we propose that distinct progenitor pools generate the posterior-most segment of the adult midgut and the HZ. This contrasts with the model that these cell populations are formed by trans-differentiation of larval hindgut cells (Takashima et al., 2013).

### Midgut stem cells adjacent to the adult HZ are less proliferative and resist tumor formation

Given the unique gene expression and lineage of the adult HZ and its juxtaposition to developmentally distinct ISC/EBs in the neighboring midgut, we next examined whether the HZ microenvironment might influence adjacent midgut cell cycle activity. A recent study (Tian et al., 2016) used *wg* pathway mutants to interrogate the function of *wg* signaling in the developing midgut. They found that *wg* signaling is important for fate establishment and proliferation control near the midgut/hindgut boundary. However, it remained unclear whether this phenotype resulted from disrupting development or adult homeostasis. Further, given our identification of distinct cell types, including the HZ at the midgut/hindgut boundary, the role of each cell population at this boundary was also unclear. We thus investigated whether inter-organ (HZ/hindgut to midgut) regulation of midgut ISCs occurs specifically during adult homeostasis and injury repair, and identified which specific cell types are involved.

We first examined homeostasis. A previous report found *frizzled-3* (*fz3*), a downstream effector of the Wg pathway, to be expressed in the vicinity of Wg boundaries and in some adult ISCs/EBs, including the most posterior midgut region (R5/P4; Tian et al., 2016). Using a *fz3* reporter, we find that a subpopulation of the R5/P4 midgut region ISC/EB cells (~20%) are *esg+,fz3*+ (Fig3A-A”). These *fz3+* midgut ISC/EB cells are primarily found within 30μm of the HZ (Fig3B). We next closely examined cell cycle dynamics in this 30μm region relative to other midgut cells that were considered to be in the same region (R5/P4; Buchon et al., 2013; Marianes and Spradling, 2013). Our results show that this region consists of two cell populations with distinctly different cell cycle rates. By feeding mature adults the thymidine analog BrdU to mark S-phase activity, we noticed a significant reduction in *esg*+ ISC/EB cell cycling within 30μm of the midgut/hindgut boundary, as defined by FasIII (avg# *esg*+BrdU+ cells: 0-30μm=2.5±1.2 vs. 70-100μm=12.3±2.6 vs. total *esg+* cells: 0-30μm=6.0±2.2 vs. 70-100μm=16.2±3.2, Fig3C-C”,D). We also noted this reduction when examining all BrdU labeled midgut cells, which includes the endocycling enterocytes (avg# BrdU+ cells: 0-30μm=7.5±2.7 vs. 70-100μm=24±3.8, SupplFig3A).

**Figure 3.**
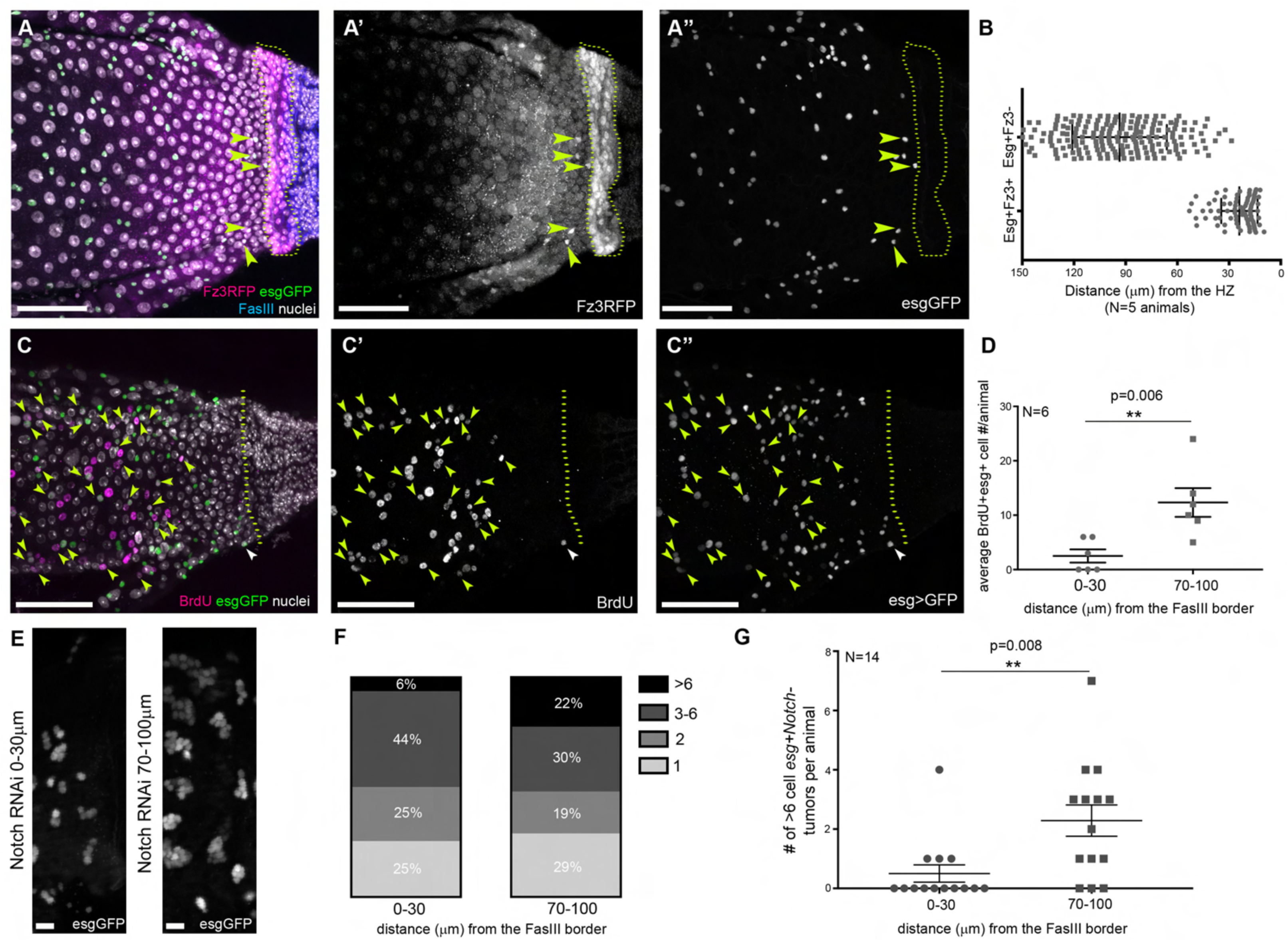
Midgut stem cells adjacent to the adult HZ are less proliferative and resist tumor formation. (A) *esg+* cells near the HZ are *fz3+*, yellow arrowheads indicate *esg+fz3+* cells, scale=50μm. (B) *esg+fz3+* are mostly within 30μm of the HZ. Data represent mean ± SEM of all *esg+fz3+* from 6 animals. (C) Near the HZ fewer cells cycle; yellow arrowheads indicate *esg+*BrdU+ cells beyond 30μm, white arrowhead indicates a *esg+*BrdU+ cell within 30μm, scale=50μm. (D) Significantly fewer *esg+* cells are BrdU+ 0-30μm from the HZ compared to 70-100μm. (E) *Notch* RNAi stem cell tumors are smaller near the HZ, scale=10μm. (F-G) *Notch* RNAi stem cell tumors of >6 cells are found less frequently 0-30μm from the HZ compared to 70-100μm. (F) Percentages of *esg+* tumor sizes in *Notch* mutant animals 0-30μm from the HZ compared to 70-100μm. Chi-Square Fisher’s Exact test, p=0.002. (G) More >6 cell *esg*+ tumors in *Notch* mutant animals are found in 0-30μm from the HZ compared to 70-100μm. Data represent mean ± SEM. Genotypes and markers indicated in panels, yellow dotted lines indicate the HZ.

As a second way of examining whether ISCs adjacent to the HZ are differentially regulated during homeostasis, we asked if the reduced cell cycle rates observed in the 30μm midgut region adjacent to the HZ impact tumor development. Previous studies in the midgut revealed that loss of *Notch* in ISCs leads to the formation of stem cell tumor-like hyperplasia (Guo and Ohlstein, 2015; Micchelli and Perrimon, 2006; Ohlstein and Spradling, 2006, 2007). We examined the posterior midgut adjacent to the HZ in animals expressing *Notch RNAi* specifically in *esg+* ISCs and associated EBs. As a proxy for tumor formation, we counted clusters of *esg*+ ISC/EBs (see Methods). Several days after *Notch RNAi* expression, within 70-100μm of the HZ we observed tumors with >6 *esg+* cells frequently (22%, see Methods). However, within 0-30μm we rarely observed these larger tumors (~6%, Fig3E-G,SupplFig3CE’), which mirrored the reduction in cell cycle rates we observed in this region (Fig3C-D). Indeed, as the distance from the HZ increases we noted an increase in the size of the tumors after loss of *Notch* in ISCs (SupplFig3B,D-E’). To confirm that this difference in *Notch* tumor formation was not due to the lack of *Notch* ligand, we examined the localization of Delta (Dl) in the midgut, and readily detected Dl+, *esg*+ ISCs/EBs throughout the posterior-most region of the midgut (SupplFig3F). In summary, during adult homeostasis, we find a unique ISC/EB cell population that lies adjacent to the HZ. Relative to neighboring midgut ISCs, which were previously considered as part of the same cell population, this population exhibits cell cycle rates that are much lower than their neighbors, are *fz3+*, and are resistant to stem cell tumor formation after loss of Notch signaling. We define these cells as organ-boundary intestinal stem cells (OB-ISCs).

### Injury to the adult HZ/hindgut drives OB-ISCs into the cell cycle

We next examined how the HZ and neighboring OB-ISCs respond to injury in adults. We previously injured the adult pylorus and identified putative stem cell activity either in or near the Wg ring (Fox and Spradling, 2009). Having now characterized at single-cell resolution the architecture and developmental origin of this region, we injured the adult HZ and hindgut and examined whether stem cell division is part of the response to injury at this organ boundary. To do this, we used *byn*Gal4 to drive the pro-apoptotic genes *head involution defective* (*hid*) and *reaper* (*rpr*) specifically in adults (see Methods). Cell death, monitored by Death caspase-1 (Dcp-1) accumulation, was only induced in the HZ and pylorus (Fig4A-C). Cell cycle re-entry after injury, as monitored by BrdU, is specific to the region surrounding the midgut/hindgut boundary (SupplFig4CvsD). As previously reported, cells in the pylorus either die or re-enter the endocycle to repair damage via cellular hypertrophy (Losick et al., 2013, Fig4D-E’). Cells in the HZ also undergo cell death, as evidenced by Dcp-1 and TUNEL staining (Fig4A-B’, SupplFig4AB’), whereas HZ cells that do not undergo cell death re-enter the cell cycle (Fig4E).

**Figure 4.**
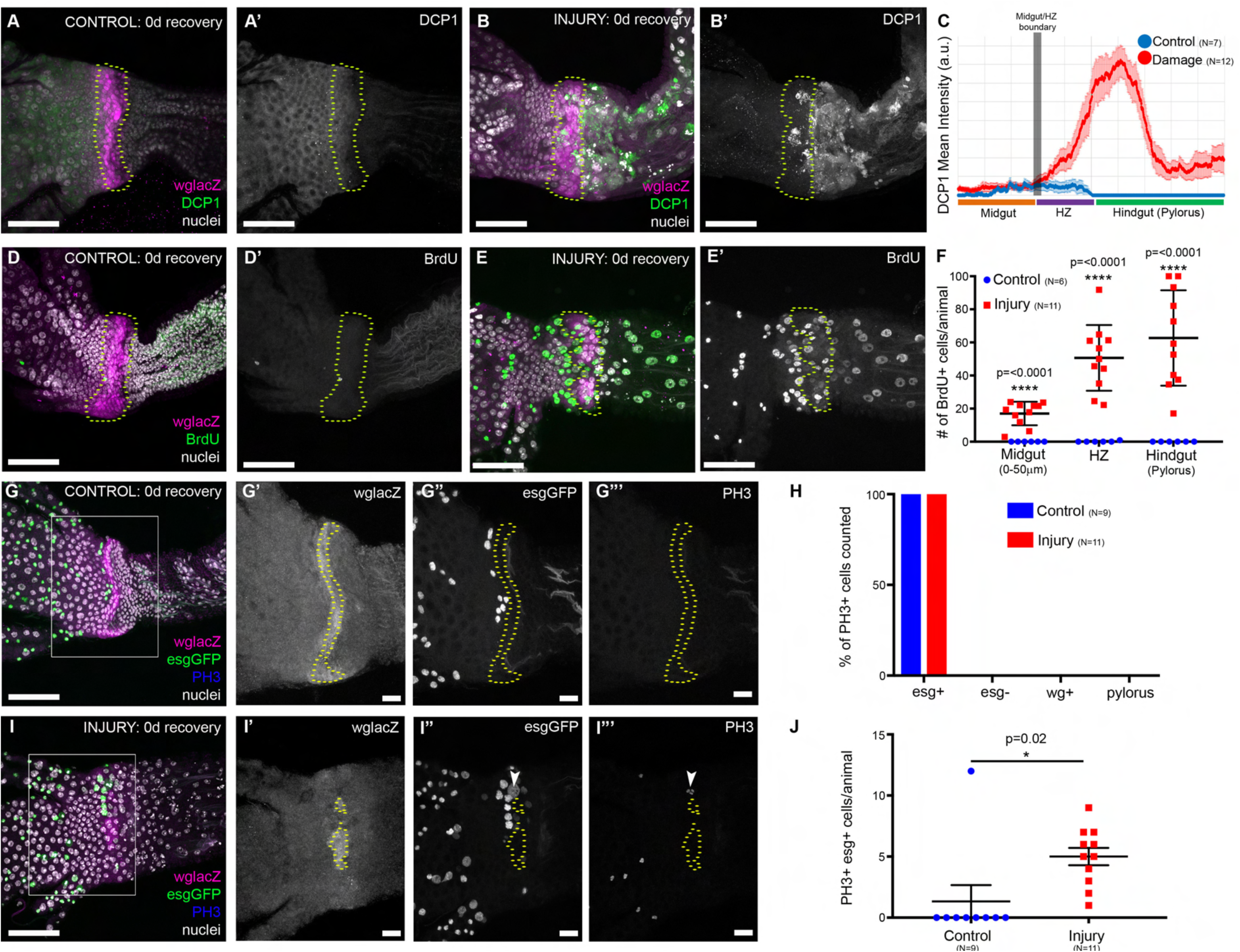
Injury to the adult HZ/hindgut drives OB-ISCs into the cell cycle. (A-B) Injury is specific to the HZ/hindgut. DCP-1 is not observed in the absence of injury (A). After injury, via *byn*Gal4 driven apoptosis, DCP1 is seen in the HZ/hindgut, but not the midgut (B), scale=50μm. (C) Line profile of relative DCP1 levels in control vs. injury. Data represent mean ± SEM. (D-F) Injury induces cell cycle re-entry. Few cells cycle in the absence of injury (D), while many cells re-enter the cell cycle after injury, even in the midgut (E), scale=50μm. (F) Quantification of BrdU+ cells in the posterior-most midgut, HZ, and the hindgut. Data represent mean ± SEM. (GJ) Injury to the HZ promotes *esg+* mitosis. *esg+*PH3+ cells are rarely observed in absence of injury (G), but are frequently observed after injury (I), scale=50μm and 10μm. (H) Only *esg+* are PH3+ with or without injury. Data represents the % of PH3+ that were also *esg+, esg-, wg+*, or FasIII+ (pyloric). (J) More cells are esg+PH3+ after injury. Data represent mean ± SEM. Genotypes and markers indicated in panels, yellow dotted lines indicate the HZ.

While cell cycle re-entry was expected in *byn+* cells, what was unexpected was our observation that *byn-* cells in the adjacent midgut also re-enter the cell cycle, despite not being injured (Fig4E-F, SupplFig4A-B). We previously found a localized population of putative injury-responsive stem cells that is positive for the mitotic marker Phospho-Histone H3 (PH3), but it remained unclear whether these cells were in the Wg ring/HZ. To clarify this, we examined PH3 along with markers for the HZ (*wg*) and OB-ISCs and associated EBs (*esg*). Strikingly, we never observed PH3+ HZ or pyloric cells, either with or without injury (Fig4G-I). These data show that the adult hindgut and associated HZ do not retain any proliferative capacity, but instead these cells undergo injury-induced endocycles (Losick et al., 2013). Further, we find all hindgut injury-induced PH3+ cells are *esg*+. After injury, *esg+* cells are more frequently PH3+ (Fig4J). We also examined Dl expression after injury, and found an increased incidence of Dl+*esg*+ cells at the midgut/hindgut border after injury, indicating that PH3+ OB-ISCs likely divide to expand after injury (SupplFig3F-G). These results show that after injury to the HZ and pylorus in adult animals: 1) the pylorus and the HZ undergo the endocycle, but do not proliferate, and 2) *esg+* cells (likely the OB-ISCs) proliferate.

### Injury to the adult HZ/hindgut causes OB-ISC hyperplasia

We next determined the long-term outcome of induced midgut OB-ISC proliferation after injury to the adult HZ/hindgut. To do so, we examined the number of OB-ISCs and their daughter EBs (*esg+*) in the range of 0-30μm and 70-100μm from the HZ after 0, 5, 10, 20, 30, and 40 days of recovery. We found that *esg+* cells close to the HZ (0-30μm) in injured adult animals expand compared to controls (Fig5A), confirming that OB-ISCs divide to expand in number after injury. Strikingly, when comparing the 0-30μm to 70-100μm interval in injured animals we find that this expansion of *esg+* cells is specific to the 0-30μm range population, especially beyond 20 days after recovery (Fig5A). This suggested that the OB-ISC response to injury, in a neighboring organ, is a locally restricted response. Using more precise bins of 10μm within a 110μm range from the HZ, we further find that the increase in *esg*+ cells originates very close to the HZ, and gradually expands in an anterior direction (SupplFig5A).

**Figure 5.**
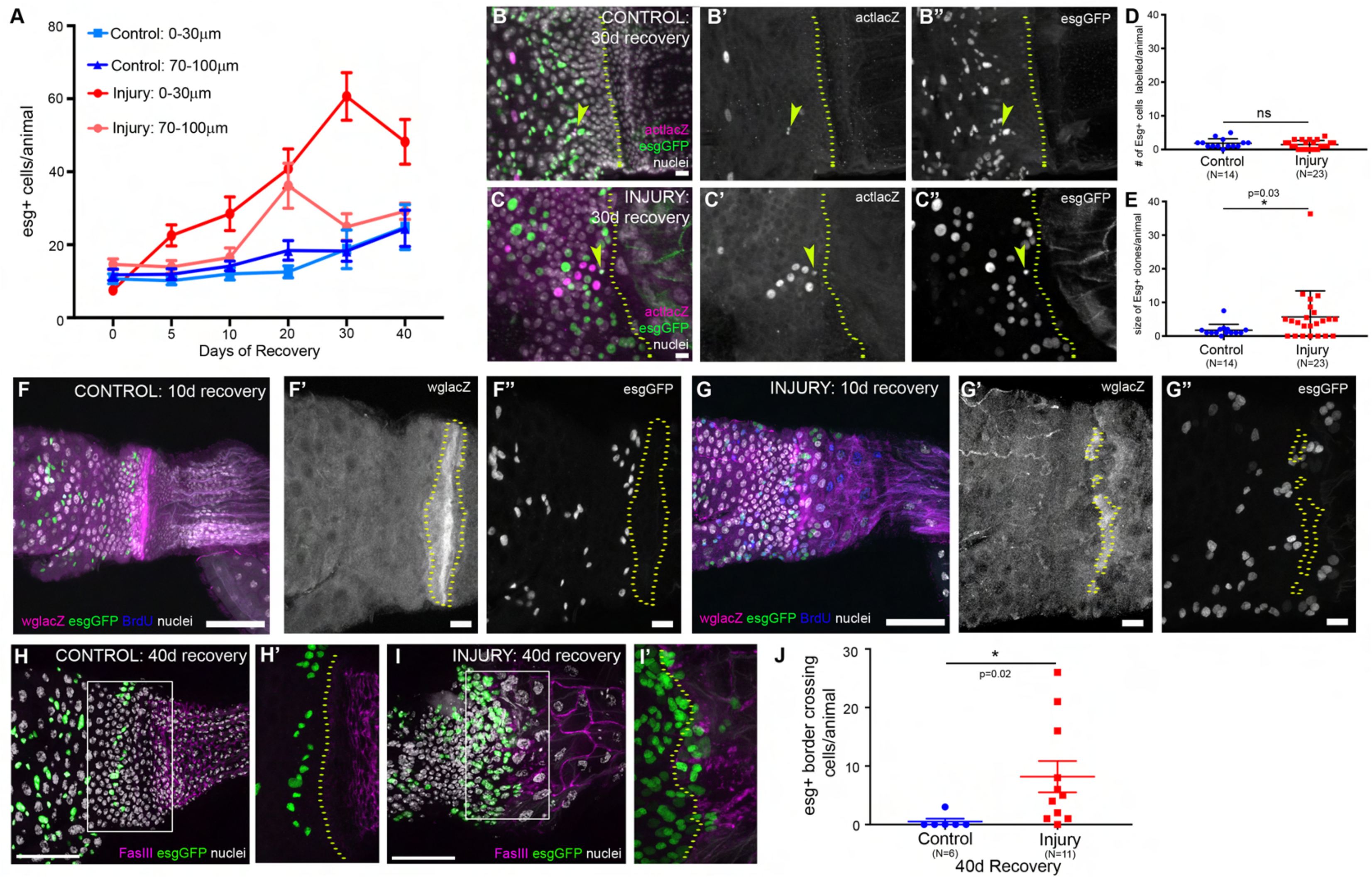
Injury to the adult HZ/hindgut causes OB-ISC hyperplasia. (A) After injury, *esg+* cells near the HZ expand. Data represent mean ± SEM. (B-E) This expansion is driven by *esg+* cell division. (B) Control and (C) Injury clone examples, yellow arrowheads indicate a marked *esg+* cell clone, scale=10μm. (D) Lineage tracing marks a similar number of *esg+* cells. Data represent mean ± SEM. (E) esg*+* clones are larger after injury. Data represent mean ± SEM. (F-G) Injury creates breaks in the HZ. In the absence of injury the HZ remains intact (F), while after injury breaks are frequently observed and *esg+* are found posterior to the HZ (G),scale=50μm and 10μm. (H-J) *esg+* cells cross the FasIII boundary. *esg+* cells rarely cross the boundary in the absence of injury (H). After injury, *esg+* cells do cross the boundary (I), scale=50μm. (J) Quantification of *esg+* cells that cross the boundary after injury. Data represent mean ± SEM. Genotypes and markers indicated in panels, yellow dotted lines indicate the HZ.

We next considered two possibilities for the expansion of OB-ISCs near the injured HZ. The first possibility was that this expansion represents symmetric ISC division, potentially reflecting an inability of OB-ISCs to differentiate (Antonello et al., 2015; Chen et al., 2016; Meng and Biteau, 2015; O’Brien et al., 2011; Zhai et al., 2015). The second possibility was that OB-ISCs both expand in number and generate new enterocytes, perhaps in response to the loss of the HZ after injury (symmetric and asymmetric divisions). We distinguished between these possibilities by combining FLP-mediated lineage tracing and Gal4-mediated genetic cell ablation injury systems (each described earlier) in an *esg-GFP* animal. A similar number of *esg+* cells are labeled per animal in both control and injury conditions (1.9±0.4 vs 1.4±.3,Fig5D). However, *esg+* clones are larger in injured animals, supporting the idea OB-ISCs divide more frequently after injury (1.7±0.5 vs. 5.7±1.6,Fig5B-C,E). *esg-* cells are labeled at the same frequency in control and injured animals (7.6±1.2 vs. 5.2±1.0,SupplFig5B) but clones are also of a similar small size (nearly always 1 cell, occasionally 2 cells), as expected given that ISCs are the source of the vast majority of cell division in the midgut, with or without injury (1.6±0.2 vs. 1.1±0.1,SupplFig5C). Of the multicellular *esg+* cell containing clones, almost all clones examined also contained *esg-* cells, supporting the model that injury promotes both symmetric and asymmetric divisions of OB-ISCs (SupplFig5D). This may reflect an inter-organ tissue repair response (see discussion).

A likely role of an organ boundary is to prevent mixing of cells from distinct organs. To examine this possibility, we tracked *wglacZ* expression following injury to the adult HZ/hindgut and recovery to observe the morphology of the HZ. In animals where injury was more severe, at 10 days of recovery we find frequent breaks in the Wg region (average breaks per animal: 2, average length of break: 12μm, Fig5F-G, SupplFig5E-F). Given that we observe gaps in the HZ, we next examined whether the excess OB-ISCs entered the hindgut through these gaps. We find that *esg*+ cells often invade breaks in the HZ 10d after injury (Fig5F”vs5G”). Indeed, more breaks in the HZ leads to more OB-ISCs invading through and around the HZ (SupplFig5G). Forty days after injury we find many OB-ISCs cells that have crossed into the hindgut (Fig5H-J). Taken together, our data demonstrates that injury to the adult HZ stimulates OB-ISC division. Under severe injury conditions, the HZ is disrupted, which allows OB-ISCs to invade across the organ boundary. Our results underscore the role of the adult HZ in repressing division and maintaining organ identity on both sides of the organ boundary.

### OB-ISCs respond to injury-induced Upd3

Given the gradual increase in OB-ISC proliferation next to the injured adult HZ, we next searched for molecular mechanisms driving this hyperplasia. The JAK-STAT ligand *upd3* (IL-6-like) is known to promote the proliferation of ISCs after enterocytes loss (Biteau et al., 2008; Jiang et al., 2009). We find that *upd3* is robustly expressed in the HZ and pylorus after injury (Fig6AvsB). Next, we used *wg*Gal4 to express ectopic *upd3* specifically in the adult HZ and find that this is sufficient to drive neighboring midgut cells to re-enter the cycle, as measured by BrdU incorporation (Fig6C-E). This response is largely limited to the midgut. BrdU+ pyloric cells are occasionally observed after *upd3* overexpression, but are only found within 5-10μm of the HZ, while BrdU+ midgut cells are frequently found ~100μm away from the HZ (Fig6D, BrdU+ pyloric cells: control=0.3±0.3, injury=6.6±2.8). This suggests that HZ-derived *upd3* moves uni-directionally towards the OB-ISCs, possibly reflecting polarized JAK-STAT signaling (Sotillos et al., 2008). Next, to test whether pylorus-derived *upd3* is sufficient to drive pyloric cells into S-phase, we drove ectopic *upd3* expression in the adult hindgut, using *byn*Gal4. Unlike HZ-specific *upd3* expression, in *byn-*induced *upd3-*expressing animals we observe a robust pyloric cell cycle re-entry response (SupplFig6AvsB). To confirm that ectopic *upd3* promotes a similar cell cycle response (i.e., mitosis) as seen in OB-ISCs after injury, we examined the mitosis marker PH3 and found that ectopic *upd3* expression from the HZ is sufficient to increase proliferation adjacent to the HZ (Fig6F-H). Given that we previously found that only ISCs divide with or without injury (Fig4H), we conclude that *upd3* expression is sufficient to promote OB-ISCs proliferation. Taken together, our data suggests that injury to the adult HZ both 1) relieves the proliferation block of OB-ISCs near the HZ and 2) promotes proliferation of OB-ISCs by inducing JAK-STAT signaling.

**Figure 6.**
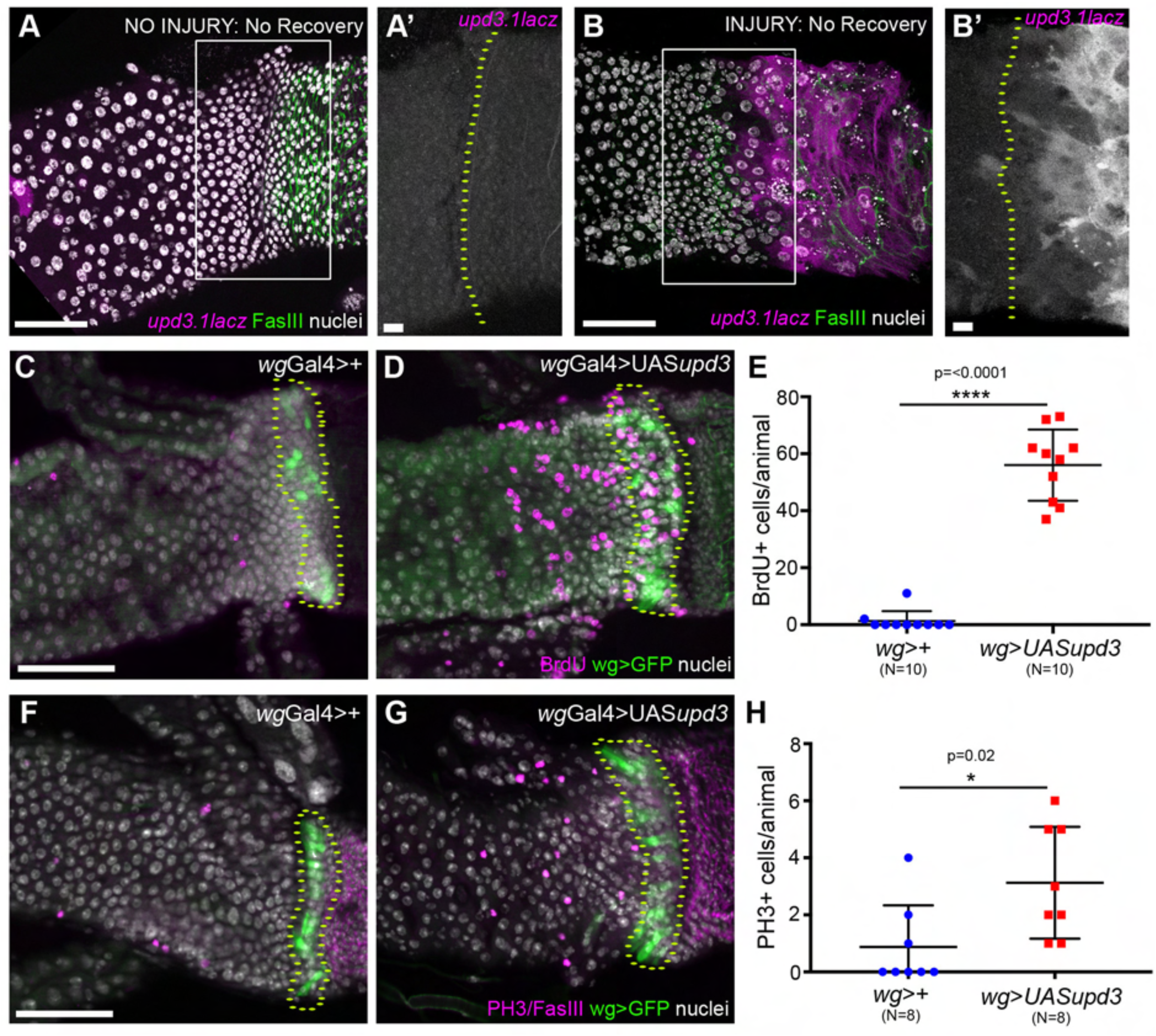
OB-ISCs respond to injury-induced Upd3. (A-B) *upd3* is induced after injury. *upd3* is rarely observed in the absence of injury (A), but is strongly induced after injury (B), scale=50μm and 10μm. (C-D) Ectopic *upd3* expression, with *wg*Gal4, drives cells in the HZ and midgut cells to re-enter the cell cycle, scale=50μm. (E) The number of BrdU+ cells in the HZ and midgut after injury is significantly more than without injury. Data represent mean ± SEM. (F-G) Ectopic upd3 expression results in more PH3+ cells in the midgut, scale=50μm. (H) The number of PH3+ cells in the midgut after injury is significantly more than without injury. Data represent mean ± SEM. Genotypes and markers indicated in panels, yellow dotted lines indicate the HZ.

## DISCUSSION

Very little is known about control of cell fate and proliferation across boundaries of functionally distinct organs. Similarly, we have a poor understanding of how stem cells in one organ are influenced by a juxtaposed organ. Our work demonstrates that a HZ, with gene expression from both the endodermal midgut and ectodermal hindgut, acts during adult homeostasis to repress normal or tumorous (*Notch* RNAi) proliferation in nearby *fz3+* OB-ISCs of the midgut (Fig7A). Injury to the adult hindgut and HZ alleviates this proliferation block, through release of *upd3* cytokine in the HZ. By examining the HZ long term after injury recovery, we observe that in cases where the HZ is significantly injured, its absence promotes invasion of hyperplastic OB-ISCs across the inter-organ boundary (Fig7B). Collectively, our results establish the midgut/hindgut boundary as a model to understand both inter-organ stem cell regulation and hyperplastic growths that invade across organ boundaries.

**Figure 7.**
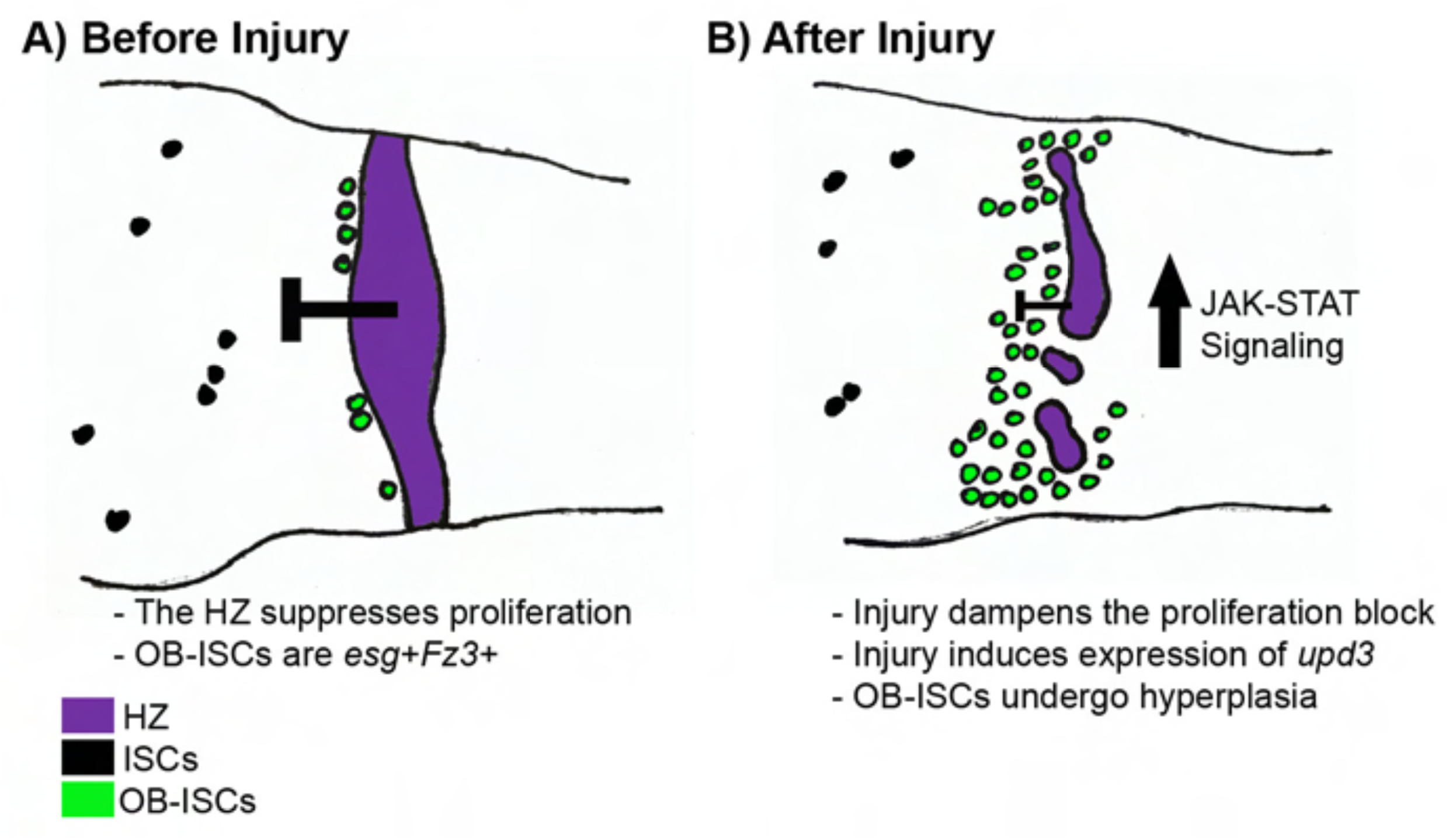
Model. (A) Before Injury: OB-ISCs reside adjacent to the HZ, which suppresses proliferation. (B) After Injury: Injury disrupts the HZ and activates JAK-STAT signaling, which promotes OB-ISCs proliferation and subsequent hyperplasia.

### The Drosophila adult hindgut lacks proliferative cells, even under injury conditions

Previously, we refuted the claim that the Wg ring/HZ harbored cells that were constitutively active hindgut stem cells, which had led to this region being termed the “hindgut proliferative zone (HPZ)” (Takashima et al., 2008). Our previous work did leave open the possibility that the Wg ring cells may serve as tissue repair stem cells, only proliferating after injury (Fox and Spradling, 2009). However, the detailed marker analysis conducted in this study revealed that it is in fact the ISCs immediately adjacent to the Wg ring/HZ, termed OB-ISCs, which proliferate in response to hindgut/HZ injury in adults. Thus, it is now clear that there is no adult HPZ, even under injury conditions. Future work can determine if asymmetric divisions of OB-ISCs are capable of re-populating the HZ by producing new HZ enterocytes, as part of an inter-organ stem cell repair response.

### Use of a hybrid zone as a strategy to maintain two distinct organs

Our study reveals that use of a HZ is a strategy for transitioning between cell fates at an organ boundary. We postulate that the HZ has two important roles in the *Drosophila* intestine: 1) to maintain organ integrity throughout development and 2) to mediate the transition between two distinct organs in both function and proliferative capacity. First, the HZ is a distinct landmark that likely directs the adoption of appropriate cell fates and coordinates morphogenesis of two structurally distinct organs during development. Second, the midgut is an actively proliferative adult tissue, while the hindgut is post-mitotic. The pylorus, analogous to the mammalian ileocecal valve, is a contractile valve that helps to move food through the intestine. Under steady-state conditions the HZ may suppress proliferation near this contractile sphincter tissue to keep the lumen clear of apoptotic cells and promote the flow of intestinal contents. Thus, accurately defining the molecular identity of intestinal boundaries will further understanding of both the development and maintenance of such boundaries.

### Organ boundary stem cells likely reside in specialized microenvironments

In *Drosophila* and mammals, the A-P pattering of the intestine is first specified by gradients of several growth factor pathways, including Wnt, and further sharpened by Hox-family homeodomain proteins and region-specific transcription factors (Nakagoshi, 2005; Zorn and Wells, 2009). In the mammalian intestine, there are both clear and imprecise morphological boundaries, but the molecular determinants of both these types of boundaries are often less clear (San Roman and Shivdasani, 2011). An example of a GI disease that develops at an intestinal boundary is Barrett’s esophagus, a metaplasia in which squamous distal esophageal cells are replaced with columnar stomach/intestine-like cells (Falk, 2002; Shaheen and Richter, 2009).

In our model, we find that after injury to the adult HZ/hindgut, OB-ISCs undergo hyperplasia and invade across an organ boundary in response to injury. This is remarkably similar to what is seen in a murine model of Barrett’s esophagus, in which a small population of residual embryonic cells that resides at the squamocolumnar junction is activated upon injury and leads to their aberrant expansion across the esophagus/stomach boundary (Wang et al., 2011). This observation, along with our current work, supports the idea that stem cell displacement, and not oncogenic mutations, may be sufficient to initiate dysplasia. Importantly, dysplasias are increasingly appreciated to set the stage for cancer progression in the intestine (Flejou, 2005; Harpaz and Polydorides, 2010).

Our work further suggests that OB-ISCs are influenced by their local microenvironments, and can respond to injury in a neighboring organ. Transcriptional profiling of the midgut has revealed that within the midgut organ, ISCs have different gene expression signatures based on their position (Dutta et al., 2015; Marianes and Spradling, 2013). Interestingly, ISCs in the most posterior portion of the midgut (R5) have a higher expression of JAK-STAT pathway genes, suggesting that these ISCs may be more poised to respond to injury induced-*upd3* that we observe in adults (Dutta et al., 2015). Indeed, a recent study in a murine model of Barrett’s esophagus suggests that intestinal injury can increase IL-6 (mammalian *upd3* homolog), which contributes to the progression of Barrett’s and subsequent development esophageal adenocarcinoma (Quante et al., 2012). Taken together, our data highlights the importance of inter-organ boundaries as specialized stem cell microenvironments, which likely strongly influences stem behavior during both homeostasis and after injury.

In conclusion, our work provides insight into how cells residing at organ boundaries function to maintain boundary integrity during adult homeostasis and after tissue injury in adult animals. Further, our results highlight the importance of future studies of stem cells at organ boundaries as they likely reside in unique microenvironments that will differentially influence their behavior.

## MATERIALS AND METHODS

### Fly stocks

All flies were raised at 25°C on standard media unless noted otherwise (Archon Scientific, Durham NC). Flybase (<u>http://flybase.org</u>) describes full genotypes for the following stocks used in this study from the Bloomington Drosophila Stock Center: *wg>lacZ* (#1567), *wg>Gal4* (#48754), *act>>lacZ* (#6355), *hsFLP^12^* (#1929). The other stocks were generous gifts: *byn>Gal4* (Singer et al., 1996), *esg>Gal4* (Bruce Edgar, University of Utah), *Myo1a>Gal4* (Ken Irvine, Rutgers), *upd3.1lacZ* and *UASupd3* (Huaqi Jiang, UTSW), UAS*Notch*RNAi (Sarah Bray, University of Cambridge), *fz3*RFP (Andrea Page-McCaw, Vanderbilt), *esg*GFP (Carnegie Fly Trap, #P01986).

### Drosophila Genetics

For developmental lineage analysis, we heat shocked (37°C water bath) *hsFLP; esgGFP; actin-FLPout-lacZ* L3 larvae for 40min, aged flies until adulthood (4d after eclosion) at 25°C. Clones were defined as any lacZ+ nuclei that were adjacent to each other (thus single labeled cells were not scored, SupplFig2B-B’). To examine *Notch* tumor-like growths, we used *esg>Gal4* to drive *UASNotchRNAi* for 10-15 days at 30°C. As a proxy for tumor growth, we counted the number of *esg+* cells, although we acknowledge that this is an under-estimate of tumor size, as *Notch* tumors are also known to contain an excess of *esg-*, Pros+ EE cells. To induce injury we aged newly eclosed flies at 18°C for 4 days, shifted to 30°C for 48hrs to induce injury, shifted to 18°C for recovery for the number of days indicated in figures (genotype: *esg*GFP/*wglacZ*; *byn>Gal4, tubGal80^ts^, UAShid, UASrpr/+* or *TM6B/+* for controls). For overexpression of *upd3*, we aged newly eclosed flies at 18°C for 4 days, shifted to 30°C for 4 day to induce *upd3* expression with either wgGal4 or bynGal4.

### Cell cycling and cell death assays

To determine cycling in WT animals, 2d old animals were fed 2.5mg/mL BrdU in 5% sucrose for 5 days. For injury BrdU feedings, flies were fed 2.5mg/mL BrdU in 5% sucrose for 24hrs on day 2 of the 30°C shift. To detect mitotic cells, flies were fed 0.2mg/mL colcemid (Sigma) in 5% sucrose for the last 12hrs of the 2 day 30°C shift. TUNEL was performed with the in situ cell death detection kit (Roche, Basel, Switzerland) as described previously (Schoenfelder et al., 2014).

### Fixed Imaging

Electron microscopy was performed as described in (Schoenfelder et al., 2014). Tissues were dissected in 1X PBS and immediately fixed in 1XPBS, 3.7% paraformaldehyde, 0.3% Triton-X for 30min. Immunostaining was performed in 1XPBS, 0.3% Triton-X, and 1% normal goat serum. The following antibodies were used in this study: FasIII (DSHB, 7G10, 1:50), Beta-Galactosidase (Abcam, ab9361, 1:1000), Prospero (DSHB, MR1A, 1:20), DCP1 (Cell Signaling, Asp261, 1:1000), BrdU (Serotec, 3J9, 1:200), Phospho-Histone 3 (Cell Signaling, #9706, 1:1000), Delta (DSHB, C594, 1:50), Pdm1 (generously shared by Steve Cohen, 1:10). All secondary antibodies used were Alexa Fluor dyes (Invitrogen, 1:500). Tissues were mounted in Vectashield (Vector Laboratories Inc.). Images were acquired with the following: an upright Zeiss AxioImager M.2 with Apotome processing (10X NA 0.3 EC Plan-Neofluar air lens or 20X NA 0.5 EC Plan-Neofluar air lens), an inverted Zeiss LSM510 (10X NA 0.3 EC Plan-Neofluar air lens or 40X NA 1.3 EC Plan-Neofluar oil immersion lens) or an inverted Leica SP5 (40X NA 1.25 HCX PL APO oil immersion).

### Image analysis

Image analysis was performed using ImageJ (Schneider et al., 2012), including adjusting brightness/contract, Z projections, and cell counts. Line prolife analysis (Fig1) was performed as follows: For each animal, two 150μm rectangular areas were selected, spanning the region from the center of Wg ring into 1) the midgut [-150 to 0μm] or 2) the pylorus [0 to 150μm]. An additional area with no visible fluorescence was selected to calculate threshold intensity per channel per animal. Each rectangular selection was then broken up to approximately 500 lines, and each line was analyzed separately for RGB intensities per pixel (0.21μm) using a publicly available RGB_Profiler ImageJ plugin (Laummonerie and Mutterer, Strasbourg, France). For each animal, the analysis produced ~350,000 individual measurements per channel per rectangle. Using R (3.3.1), the values were then averaged across the 500 lines to produce mean intensity/distance from the Wg ring. Relative fluorescence intensity was calculated by subtracting the threshold intensity per animal from the mean pixel intensity, producing a single dataset containing the relative fluorescence intensity means for each pixel per animal (~1500 data points per animal). Last, the relative fluorescence intensity was averaged across animals, producing a single relative fluorescence intensity per pixel across a 200μm distance.

## Acknowledgments

The following kindly provided reagents used in this study: The Bloomington *Drosophila* Stock Center, Developmental Studies Hybridoma Bank, Ken Irvine, Sara Bray, Bruce Edgar, Huaqi Jiang, Andrea Page-McCaw, and Steve Cohen. Ruth Montague, Emily Bowie, Michael Sepanski, and Benjamin Stormo provided valuable technical assistance. We thank Ken Poss, Purushothama Rao Tata, Jacob Sawyer, and Fox laboratory members for valuable comments on the manuscript. This project was supported by NIGMS grants GM118447 to DF and GM109538 to JS, as well as a Duke Regeneration Next Initiative postdoctoral fellowship to JS.

